# Lysyl Oxidases are Necessary for Myometrial Contractility and On-Time Parturition in Mice

**DOI:** 10.1101/2024.09.05.610344

**Authors:** Alexis Ouellette, Christina Do, Sydney Cohn-Guthrie, Ying-Wai Lam, Mala Mahendroo, Shanmugasundaram Nallasamy

**Affiliations:** Department of Obstetrics, Gynecology, and Reproductive Sciences, University of Vermont Larner College of Medicine, Burlington, VT, 05405, USA; Vermont Biomedical Research Network Proteomics Facility, University of Vermont, Burlington, VT, 05405, USA; Department of Biology, University of Vermont, Burlington, VT, 05405, USA; Department of Obstetrics and Gynecology, University of Texas Southwestern Medical Center, Dallas, TX 75390, USA; Cecil H. and Ida Green Center for Reproductive Biology Sciences, University of Texas Southwestern Medical Center, Dallas, TX 75390, USA

**Keywords:** Extracellular matrix, Lysyl oxidase, Myometrium, Pregnancy, Parturition

## Abstract

The extracellular matrix (ECM) plays a pivotal role in the maintenance of tissue mechanical homeostasis. Collagens and elastic fibers are the most predominant fibrous ECM proteins providing tissue mechanical function through covalent cross-linking which is mediated by the lysyl oxidase family of enzymes. In this study, the function of lysyl oxidases in maintaining the integrity of the extracellular matrix in the myometrium and its impact on parturition-timing was investigated. Gene and protein expression analyses demonstrate that a sub-set of the lysyl oxidase family of enzymes are highly induced in pregnant myometrium. Inhibition of the activity of the lysyl oxidase family of enzymes through β-aminopropionitrile (BAPN) delays parturition in mice, in part, due to myometrial dysfunction. In BAPN treated mice, the expression of genes encoding contraction associated proteins such as connexin 43, oxytocin receptor and prostaglandin synthase 2 is significantly reduced in the myometrium compared to the untreated control mice. Proteomic analysis revealed that the composition of the ECM is altered in response to BAPN treatment which demonstrates that the inhibition of the activity of lysyl oxidases disrupted the integrity of the myometrial ECM. Our findings demonstrate that the lysyl oxidases-mediated ECM function is necessary for the myometrium to transition from a quiescent to a contractile phenotype at term for on-time parturition.

## INTRODUCTION

Preterm birth (PTB) is the primary cause of perinatal infant mortality. The rate of PTB ranges from 4-16% globally with an estimated 13.4 million preterm babies born in 2020 (1). In the United States, the rate of PTB in 2022 was 10.4% which is disturbingly high for a developed nation (2,3). While the etiology of PTB is complex and multifactorial, premature myometrial contraction is a commonly observed clinical sign preceding preterm labor (4,5). Thus, understanding the mechanisms that regulate myometrial quiescence, and contractility will help us identify the risk factors and potential therapeutic targets to prevent premature myometrial contraction and its contribution to PTB. Our understanding of the mechanisms by which myometrium transitions from quiescence to contractility has improved over the decades including but not limited to fetal, hormonal, and inflammatory signaling pathways (6–9). However, an understanding of the role of biomechanical signals in this process is still evolving.

Biomechanical signals are predominantly elicited by alterations to the structure and composition of extracellular matrix (ECM). In addition to its traditional role of providing tissue architectural and mechanical properties, the ECM is involved in the regulation of multiple cellular processes such as cell proliferation, differentiation, contractility, migration, and survival (10,11). The myometrium experiences progressive mechanical loading while supporting pregnancy (12). To endure these biomechanical forces, the myometrial smooth muscle cells and fibroblasts presumably synthesize and remodel ECM. Among all known ECM components, collagen and elastic fibers are the predominant fibrous proteins known to provide tissue strength and resilience, respectively (13,14). The levels of collagen and elastin are increased in myometrium during pregnancy in mice, rats and humans (15–18). We have already demonstrated that the collagen and elastic fibers undergo structural reorganization in mouse myometrium from early to late pregnancy indicating changes to synthesis, processing and assembly (15).

The ultimate strength of collagen and elastic fibrous proteins is determined by the covalent crosslinking between lysine residues mediated by the lysyl oxidase family of enzymes (19–21). This enzyme family consists of five genes: Lysyl oxidase (LOX) and LOX like 1-4 (LOXL1-4). They encode enzymes with identical functions but exhibit tissue specific spatiotemporal expression patterns and functions. Their role in ECM homeostasis is revealed by distinct phenotypes in knockout mouse models and through their aberrant expression and activity in disease conditions including fibrosis, tumor metastasis, pelvic organ prolapse and endometriosis (19,22–24). Deletion of LOX causes aortic aneurysms and perinatal death in mice (25). Mice lacking LOXL1 are viable but exhibit pelvic organ prolapse in adult life (23). While deletion of LOXL2 or LOXL3 leads to perinatal lethality, the deletion of both LOXL2/LOXL3 results in embryonic lethality (26–28). LOXL4 deleted mice are viable (29). On the other hand, increased activity of lysyl oxidase enzymes leads to collagen crosslinking, and subsequent stiffening of the ECM which are major drivers of fibrosis, tumor progression and metastasis (19,24,30). Inhibition of LOX activity through β-aminopropionitrile (BAPN) – a natural and irreversible inhibitor of activity of all lysyl oxidases – blocks collagen crosslinking and reduces tumor progression (19,30).

Lysyl oxidases are widely expressed in rodent and human reproductive tissues including the uterus, placenta, fetal membranes, cervix, and vagina (22,23,31–35). LOX is transcriptionally regulated by steroid hormones. Estrogen induces LOX in the cervix and vagina (33,35). The estrogen induced LOX is downregulated by progesterone in the vagina (35). In this study, we have analyzed the expression profile and activity of lysyl oxidases in the mouse myometrium during pregnancy. Consistent with elevated expression and activity of lysyl oxidase enzymes in mouse myometrium, inhibition of their activity delayed parturition in mice by altering ECM composition.

## MATERIALS AND METHODS

### Animals

CD1 timed-pregnant mice used in the study were purchased from Charles River Laboratories. Mice strain C57B6/129sv mixed strain mice were maintained in a barrier facility at the University of Vermont under a 12-hour light/dark cycle. The mice used in the study were 2 to 4 months of age and nulliparous. Timed mating was performed to collect tissues at different days of pregnancy. Breeding pairs were set up in the morning and checked for the presence of a vaginal plug in the afternoon. The presence of a vaginal plug was considered day 0 of pregnancy. For BAPN experiment, the C57B6/129sv mixed strain mice were treated with a mixture of 0.8% BAPN and 5% sucrose in drinking water on the morning of gestation day 12 of pregnancy and maintained the BAPN water supply until the end of experiments. All animal studies were conducted in accordance with the National Institutes of Health Guide for the Care and Use of Laboratory Animals humane animal care standards. The Institutional Animal Care and Use Committee at the University of Vermont approved all animal experimental protocols used in this study.

### Ovariectomized mouse model for hormonal administration

Ovariectomies were performed on female CD1 mice that were 7 to 8 weeks of age. These mice were given 2 weeks of rest following the surgery. After 2 weeks, the ovariectomized mice were administered a vehicle (corn oil) or estrogen (100 ng, s/c); tissues were collected after 6 h, 12 h, 18 h or 24 h. In a second experiment, the ovariectomized mice were treated with progesterone as follows: Vehicle, P (1 mg), P (2 mg), P (1 mg) + E (10 ng), or E (100 ng) then after 2 days P (1 mg). Tissues were collected after 24 h.

### Tissue collection

Myometrial tissues were collected from non-pregnant, pregnant (at various gestational time points), and one day postpartum mice. Non-pregnant tissue was collected during the metestrus stage of the estrous cycle determined by analysis of a vaginal smear. Gestational time points were determined using a timed mating procedure as described above. All mice were euthanized using isoflurane overdose followed by cervical dislocation. Myometrial tissues were collected under a dissecting microscope, using a scalpel to scrape the endometrium off the uterus. Tissues were flash frozen in liquid nitrogen and stored at -80°C until further processing. The uterine segments along with fetal sac units were immersed in OCT (Tissue-Tek) and solidified with liquid nitrogen. The OCT blocks were then cut using a cryostat to create 5 µm tissue sections which were mounted on glass slides and stored at -80°C until further use.

### RNA isolation and quantitative polymerase chain reaction

Total RNA from frozen myometrial tissues was extracted using RNA STAT-60 (Tel-Test B) according to manufacturer’s protocol. The total RNA was then treated with DNase I (Ambion) to remove genomic DNA. The iScript reverse transcription supermix (Bio-Rad Laboratories) was used to prepare complementary DNA. Quantitative polymerase chain reaction was performed using SYBR Green and respective primers designed for our genes of interest. Gene expression was calculated according to the 2^−ΔΔCt^ method. The target gene expression was normalized to the expression of a housekeeping gene, *Rplp0*. The primers used in this study are listed in **Table 1**.

**Table 1.**
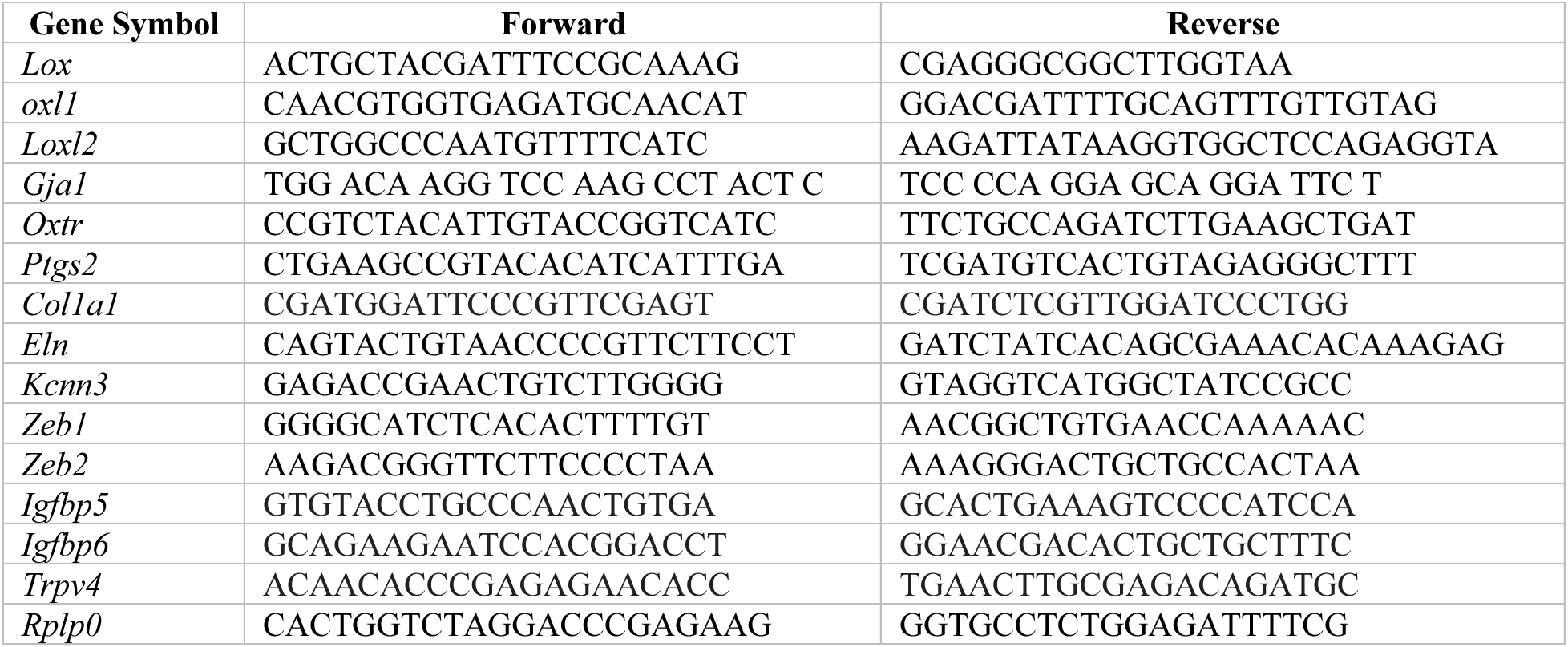
Primers used for qPCR analysis.

### Preparation of tissue lysates from myometrial tissues

*RIPA extracts:* Frozen myometrial tissues of specific gestational time points were pulverized, suspended in RIPA containing 1% protease inhibitors and EDTA (Thermo Fisher Scientific), and homogenized. The samples were centrifuged at 4°C at 13,000 rpm for 15 min to collect the supernatant protein samples. Protein concentration of the samples was estimated using a BCA protein assay (Thermo Fisher Scientific).

*PBS and urea extracts:* Frozen myometrial tissues of specific gestational time points were pulverized, suspended in phosphate-buffered saline (PBS) containing 1% protease inhibitors and EDTA (Thermo Fisher Scientific), and homogenized. The samples were centrifuged at 4°C at 13,000 rpm for 15 min. The supernatant was collected and labeled as PBS soluble protein fraction. The remaining pellet was resuspended in 6 M urea buffer containing 1% protease inhibitors and EDTA (Thermo Fisher Scientific) and homogenized. The samples were then left on a gentle rotation at 4°C for 24 h. The next day the samples were centrifuged at 4°C at 13,000 rpm for 10 min. The supernatant was collected and labeled as the urea soluble fraction. Both PBS and urea extracted sample fractions were exchanged with equivalent volume of 0.5 M Tris/0.5% SDS, pH 8.0 buffer and concentrated using Vivaspin™ ultrafiltration spin columns (Vivaspin 2, 5 kDa MWCO, # 28932359, Cytiva Life Sciences). Protein concentration of the samples was estimated using a BCA protein assay (Thermo Fisher Scientific).

### Western blot

Twenty micrograms of each protein sample in a 20:1 dilution of 4X Laemmli Sample Buffer (Bio-Rad): β-mercaptoethanol (Sigma) was boiled at 95°C for 10min. The samples were loaded, alongside a protein standard (Precision Plus Protein Kaleidoscope, Bio-Rad), into a handmade 10% Tris-HCl gel. Gel electrophoresis was run at 50 V, until the samples passed through the stacking gel, where the voltage was then increased to 100 V and run for another hour. Proteins were transferred onto a nitrocellulose membrane (Bio-Rad) at 100 V at 4°C for 1 h. Membranes were blocked in 3% nonfat dry milk in TBST (Blotting-grade blocker nonfat dry milk; Bio-Rad) at room temperature for 1h. The membranes were incubated with a primary antibody in blocking solution at 4°C overnight. The following day, the membranes were washed with TBST and incubated with their respective HRP-labelled secondary antibody at room temperature for 1 h. The membranes were imaged with equal volume of Amersham^TM^ ECL^™^ Western Blotting Detection Reagents (Cytiva Life Sciences). The primary antibodies used were: anti-mouse collagen type Ⅰ (Millipore Cat# AB765P, RRID:AB_92259), collagen type Ⅲ (Proteintech Cat# 22734-1-AP, RRID:AB_2879158), tropoelastin (Elastin Products Company Cat# PR385, RRID:AB_3099530), LOX (Abcam Cat# ab174316, RRID:AB_2630343), LOXL1 (Thermo Fisher Scientific Cat# PA5-87701, RRID:AB_2804357), LOXL2 (Novus Cat# NBP1-32954, RRID:AB_2135089), Connexin 43 (Cell Signaling Technology Cat# 3512 (also Used By NYUIHC-970), RRID:AB_2294590), and β-Actin (Cell Signaling Technology Cat# 3700 (also 3700P, 3700S), RRID:AB_2242334). The secondary antibodies used were anti-mouse IgG, HRP-linked antibody (Cell Signaling Technology Cat# 7076 (also 7076S, 7076V, 7076P2), RRID:AB_330924), anti-rabbit IgG, HRP-linked antibody (Cell Signaling, 7074S; ), goat anti-rabbit IgG (H/L):HRP (Bio-Rad Cat# STAR208P, RRID:AB_3099534) or goat anti-mouse IgG (H/L):HRP (Bio-Rad Cat# 0300-0108P, RRID:AB_808614).

### Confocal microscopy imaging

Frozen uterine tissue slides were fixed in cold acetone for 10 min, air dried for 20 min, and washed with PBS. The sections were blocked with 10% normal goat serum (Life technologies) at room temperature in a moist chamber for 30 min. Primary antibodies in normal goat serum was added to each section and incubated at 4°C overnight. The next day, the sections were washed with PBS and then incubated with Alexa Fluor conjugated secondary antibodies (Thermo Fisher Scientific) in normal goat serum at room temperature for 30 min. Slides were washed with PBS and coverslips were mounted using ProLong Gold Antifade Mountant with DNA Stain DAPI (Thermo Fisher Scientific). Images were captured using a Nikon A1R confocal microscope galvanometer scanner. The settings of the galvanometer scanner used were as follows: single illumination point scan as fast as 8 frames per second for a 512 x 512-pixel field, and up to 4096 x 4096 pixels image capture. The primary antibodies and subsequent secondary antibodies used were α-actin (1A4) (Santa Cruz Biotechnology Cat# sc-32251, RRID:AB_262054), LOX (Abcam Cat# ab174316, RRID:AB_2630343), LOXL1 (Thermo Fisher Scientific Cat# PA5-87701, RRID:AB_2804357), LOXL2 (Abcam Cat# ab96233, RRID:AB_10677617), tropoelastin (Elastin Products Company Cat# PR385, RRID:AB_3099530), Alexa-Fluor 488 goat anti-mouse IgG(H+L) (Thermo Fisher Scientific Cat# A-11029, RRID:AB_2534088) and goat anti-rabbit IgG(H+L) Alexa-Fluor 555 (Thermo Fisher Scientific Cat# A32732, RRID:AB_2633281).

### *In situ* activity assay for lysyl oxidases

The activity of lysyl oxidases was examined using an *in situ* activity assay as described (36). Briefly, the frozen tissue sections prepared from mouse uterus were fixed with neutral buffered formalin for 10 min. The tissue sections were blocked with normal goat serum for 30 min in a moist chamber at room temperature. They were incubated with 200 μM EZ-Link™ Hydrazide-Biotin (BHZ) (Thermo Fisher Scientific, Cat#: 21339) overnight at 37°C. After removing excess BHZ by washing with PBS, the sections were incubated with Streptavidin-Alexa Fluor™ 555 conjugate for 1 h in dark moist chamber. The sections were mounted with ProLong Gold Antifade Mountant with DNA Stain DAPI and imaged using Nikon A1R-ER Confocal Microscope with galvanometer scanner at 40X. The image settings were set for gestation day 15 sample, and the same settings were used for all other sections to compare staining intensity.

### Proteomics

#### Sample preparation and trypsin digestion

Urea extracted protein samples were used in this experiment. Total protein extract (50 µg) from each sample was loaded onto the SDS-PAGE and run slightly (∼ 5 mm) into the separating gel which was stained with Bio-Safe™ Coomassie G-250. SDS-PAGE allowed the assessment of protein loading and removal of any incompatible detergents for subsequent in-gel trypsin digestion. Single wide bands containing unseparated proteins from individual samples were excised, minced to approximately 1 mm^3^ cubes, followed by destaining with 50% acetonitrile (CH3CN)/50 mM triethyl ammonium bicarbonate. After dehydrating with CH3CN, reduction and alkylation of disulfides were conducted with 10 mM dithiothreitol and 55 mM iodoacetamide in 100 mM triethyl ammonium bicarbonate, respectively. After repeated washing and dehydrating with triethyl ammonium bicarbonate and CH3CN, the gel pieces were swelled in ∼12 ng/µL trypsin solution (Promega) and incubated overnight at 37°C. Peptides were extracted successively with formic acid (FA) and acetonitrile and dried under vacuum.

#### Peptide labeling by Tandem Mass Tag (TMT) and high pH fractionation

The labeling procedures were performed according to the manufacturer’s protocols (Thermo Fisher Scientific). Briefly, 50 µg of the dried peptides from each sample were resuspended in 100 µL of 100 mM triethyl ammonium bicarbonate, and 0.8 mg of TMT reagents dissolved in 41 µL of anhydrous CH3CN was added, followed by briefly vortexing and incubating for 1 h at room temperature. The reactions were then quenched by adding 8 µL of 5% hydroxylamine. Short mass spectrometry runs, and database searches confirmed that the labeling efficiency on average across all samples was > 97%, verifying the number of peptides labeled in individual samples before combining. The combined labeled peptides (∼33 µg) were fractionated using the high-pH reversed-phase spin column (Cat. No.: 84868; Thermo Scientific) into 16 fractions (10%, 13%-25% with 1% increments, 35% and 50%). All fractions were dried under vacuum and kept at -80°C until mass spectrometry analysis.

#### Liquid chromatography-tandem mass spectrometry (LC-MS/MS)

The fractionated TMT-labeled peptides were resuspended in 10 µL 2.5% CH3CN and 2.5% FA in water. Half of the labeled peptides in each fraction were analyzed. Mass spectrometry was performed using an Orbitrap Eclipse mass spectrometer coupled to an EASY-nLC 1200 system (Thermo Scientific). Samples were loaded onto a 100 μm x 350 mm capillary column packed with UChrom C18 material (1.8-µm 120 Å Uchrom C18, nanoLC-MS Solutions) at a flow rate of 300 nl min^-1^. The column end was laser pulled to a ∼3 μm orifice and packed with minimal amount of 5um Magic C18AQ before packing with the 1.8-μm particles. Peptides were separated using a gradient of 4-20% CH3CN/0.1% FA over 110 min, 20-32% CH3CN/0.1% FA in 25 min, 32-76% CH3CN/0.1% FA in 25 min then 76% CH3CN/0.1% FA for 10 min, followed by an immediate return to 4% CH3CN/0.1% FA and a hold at 4% CH3CN/0.1% FA. Peptides were introduced into the mass spectrometer via a Nanospray Flex Ion Source with a spray voltage of 1.9 kV, an Ion Transfer Tube Temperature of 275°C, and the RF set at 30%. “Cycle time in 3 seconds” acquisition mode, TMT-specific Real-Time Search (RTS) and Synchronous Precursor Selection (SPS) workflows were used to acquire mass spectrometry data. Survey scans from *m/z* 400-1400 at 60,000 resolution (AGC target: standard; max IT: auto; profile mode) was acquired with data-dependent collision-induced dissociation (CID) tandem mass spectrometry (MS/MS) scans on the most abundant ions (AGC target: standard; max IT: 35 ms; ion trap scan rate: turbo) in the ion trap with a normalized collision energy (NCE) at 35% and an isolation width of 1 *m/z*. RTS was used for triggering MS3 acquisitions with TMT6plex, carbamidomethylation, and oxidation as modifications and the Mus musculus UP000000589 FASTA as database (downloaded from Uniprot on March 29, 2021). XCorr values of 1.4 and 2 were used as score thresholds for doubly and triply charged peptides, respectively. Protein “close-out” was not enabled. MS3 was performed using SPS with isolation widths of 1 Da and 2 Da for MS1 and MS2 isolation windows, respectively, and an NCE of 65% (AGC target: custom: 200%; max IT: 200). The product ions from MS3 were scanned from *m/z* 100 – 500 in the Orbitrap with a resolution of 50,000. Dynamic exclusion was enabled (exclude isotopes: on; exclusion duration: 45 sec; charge states = 2-6). Precursor Fit window was set as 1 and the Fit Threshold as 30%. The samples were injected twice as technical replicates.

#### Database searches

The 32-mass spectrometry .RAW files (16 high-pH reversed phase separation fractions analyzed in 2 technical replicates) generated from each experiment were imported into the Proteome Discoverer 2.5 (Thermo Fisher Scientific) as “fractions”. Product ion spectra were searched using the SEQUEST in the Processing workflow against the Mus musculus UP000000589 FASTA database from Uniprot; downloaded on March 29, 2021). Search parameters were as follows: (1) full trypsin enzymatic activity; (2) maximum missed cleavages = 2; (3) minimum peptide length = 6; (4) mass tolerance at 10 ppm for precursor ions and 0.6 Da for fragment ions; (5) dynamic modification on methionine (+15.9949 Da: oxidation), dynamic modification on N-terminus (+42.01 Da: Acetyl) , static TMT6plex modification (The TMT6plex and TMT10plex have the same isobaric mass) on N-termini and lysine residues (229.163 Da); (6) 4 maximum dynamic modifications allowed per peptide; and (7) static carbamidomethylation modification on cysteines (+57.021 Da). Percolator node was included in the workflow to limit the False Discovery Rate (FDR) to less than 1% in the data set.

#### Quantification and statistical analysis

The biological replicates and technical replicates were assigned accordingly in the “Study Factors” tab (CNTRL-1, CNTRL-3, CNTRL-4, CNTRL-5, CNTRL-6, BAPN-1, BAPN-3, BAPN-4, BAPN-5, BAPN-6) and “Samples” tab, respectively in the Proteome Discoverer. The abundances of TMT-labeled peptides were quantified with the Reporter Ions Quantifier node in the Consensus workflow and parameters were set as follows: (1) both unique and razor peptides were used for quantification; (2) Reject Quan Results with Missing Channels: False; (3) Apply Quan Value Corrections: True (values set according to the product spreadsheet (Lot # XD341063)); (4) Co-Isolation Threshold: 75; (5) Average Reporter S/N Threshold = 10; SPS Mass Matches [%] = 65; (6) Normalization mode: “Total Protein Amount”; and (7) Scaling Mode was set “On All Average”. Protein ratio calculation was “Protein Abundance Based”. Two-tailed t-test was used for hypothesis testing and associated p-values were calculated for protein identities quantified across all samples. All the protein identification and quantification information (<1% FDR) was exported from the Proteome Discoverer result files to Excel spreadsheets. Fold change (log2) and p values (-log10) information and scaled abundances (the abundances were scaled so that fold changes of proteins of varying abundances could be represented on the same color intensity scale of the heat map) were imported into Graph Pad Prism 10 (GraphPad Software Inc., CA) for constructing volcano plots and heat maps.

#### Enrichment Analysis

The list of gene names of the differentially expressed proteins (fold change (BAPN/Control)>2, p<0.05) was imported into pathfindR (https://github.com/egeulgen/pathfindR) (37) with STRING as the database for the protein interaction network (PIN). The output (enriched_terms.html) is included in the supplementary table. The knitted pdf of the Rmd script that read in the relevant information from the Proteome Discoverer Excel output and ran the pathfindR is uploaded to PRIDE.

### Statistical analysis

Statistical analysis was completed using GraphPad Prism software. The student’s test was used to compare two groups. A one-way analysis of variance (ANOVA) followed by a Tukey multiple comparisons test was used to compare multiple groups. The values were expressed as the mean ± the standard error of the mean (SEM). P < 0.05 was considered as significant between groups.

## RESULTS

### LOX is significantly induced in mouse myometrium during pregnancy

To analyze the expression profile of *Lox* in mouse myometrium over the course of pregnancy, we utilized myometrial tissues collected from non-pregnant (NP) mice, at various gestational time-points starting from day 6 to day 18, and postpartum (PP) mice. The mRNA expression of *Lox* was significantly induced from gestation day 10 to term compared to NP myometrium **(Fig. 1A)**. LOX is synthesized within the cell as a pro-enzyme which is cleaved into an active enzyme after being exported into the ECM. To examine the levels of pro-LOX and active LOX in mouse myometrium, we prepared tissue lysates from the myometrial tissues by extracting with phosphate buffered saline (PBS) followed by urea to release LOX that was integrated into the ECM. The pro-LOX was easily extractable and was enriched in PBS soluble fractions. However, the active LOX was highly enriched in urea soluble fractions **(Fig. 1B and C)**. Consistent with gene expression results, the protein levels of LOX were highly elevated in the pregnant myometrium. Our subsequent immunolocalization studies revealed that LOX was predominantly localized in myometrial smooth muscle cells **(Fig. 1D)**. These results indicate that LOX is highly induced in the mouse myometrium during pregnancy and has a potential role in myometrial tissue function during pregnancy and parturition.

**Fig. 1.**
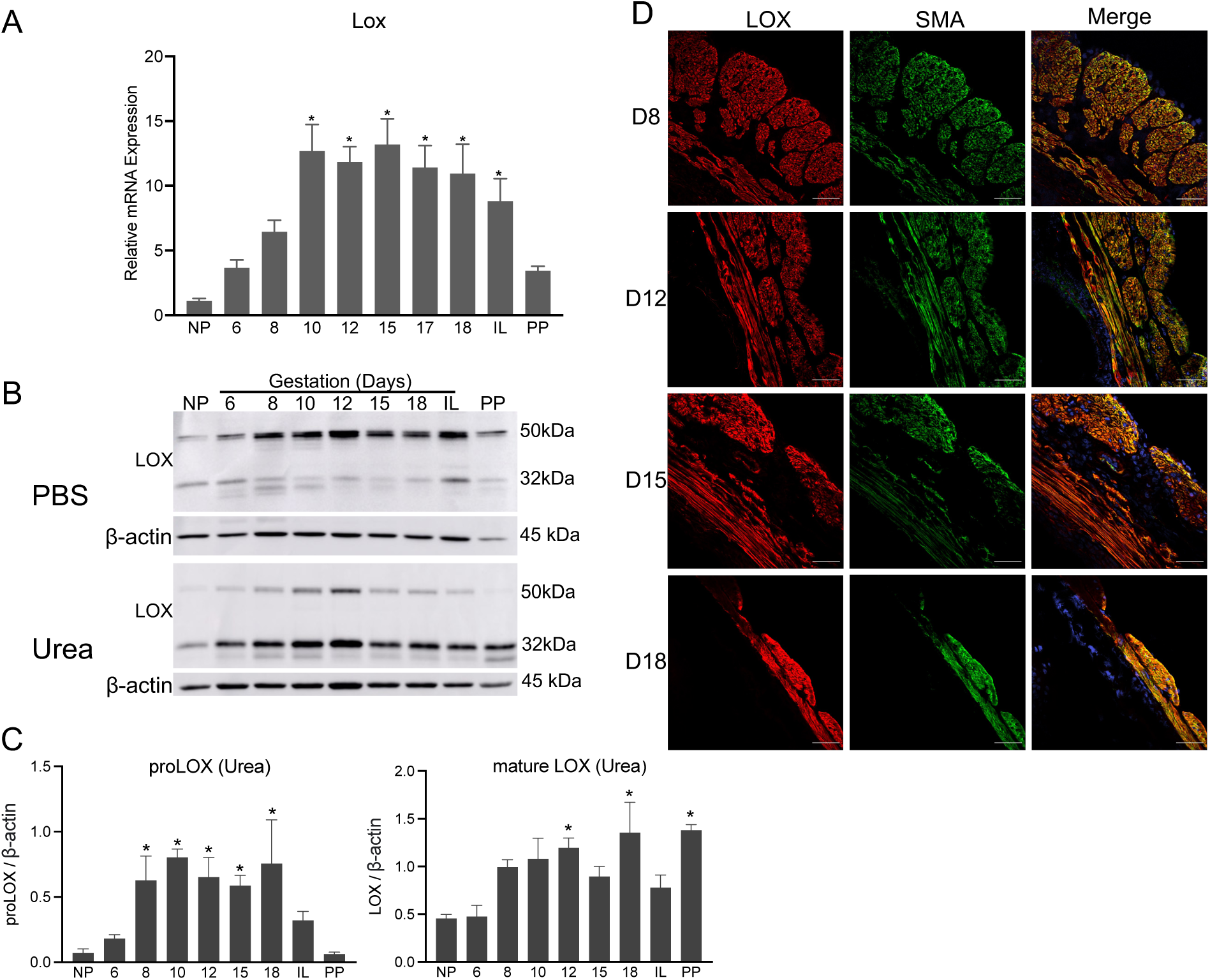
Expression and localization of LOX in mouse myometrium. **A.** Expression of *Lox* in mouse myometrium. Total RNA isolated from NP (non-pregnant), gestation day 6, 8, 10, 12, 15, 17, 18, IL (in-labor) and PP (postpartum) mice was used. The mRNA expression was normalized to *Rplp0* and compared with NP samples (n=6/group, *p<0.05). **B.** Western blot analysis of LOX protein levels in PBS and urea soluble fractions of NP, gestation day 6, 8, 10, 12, 15, 18, IL and PP mouse myometrial tissues. β-actin was used as a loading control. These are representative images from three independent replicates. **C.** Quantification of protein levels of proLOX and mature LOX in urea fractions prepared from myometrial tissues of NP, gestation day 6, 8, 10, 12, 15, 18, IL and PP mice normalized to β-actin levels and compared with NP samples (n=3/group, *p<0.05). **D.** Confocal imaging analysis of LOX in gestation day 8, 12, 15, and 18 myometrium. Representative images from three independent replicates. Brightness and contrast for each image were optimized individually for optimal visualization of morphology. Scale bar: 50 μm. LOX – Lysyl oxidase, SMA – Smooth muscle actin. Note: LOX is predominantly localized in smooth muscle tissue of the myometrium.

### LOXL1 and LOXL2 are also significantly induced in mouse myometrium during pregnancy

We next analyzed the gene expression of *Loxl1-4* in mouse myometrium. Similar to *Lox, Loxl1* and *Loxl2* were significantly induced in the pregnant myometrium **(Fig. 2A and 3A)**. The gene expression of *Loxl3* and *Loxl4* could not be detected using qPCR analysis. We next analyzed the protein levels in the whole tissue lysates prepared from myometrium. The protein levels of LOXL1 were significantly elevated in the pregnant myometrium consistent with its gene expression **(Fig. 2B and C)**. While LOXL2 protein levels appeared to be modestly elevated during pregnancy, there were high variations between replicates and thus were not statistically significant in the pregnant myometrium (**Fig. 3B and C)**. Similar to LOX, both LOXL1 and 2 were localized in the smooth muscle compartment of the mouse myometrium **(Fig. 2D and 3D)**. These results demonstrate that not only LOX, but also LOXL1-2 enzymes were significantly induced and could potentially contribute to the function of myometrium.

**Fig. 2.**
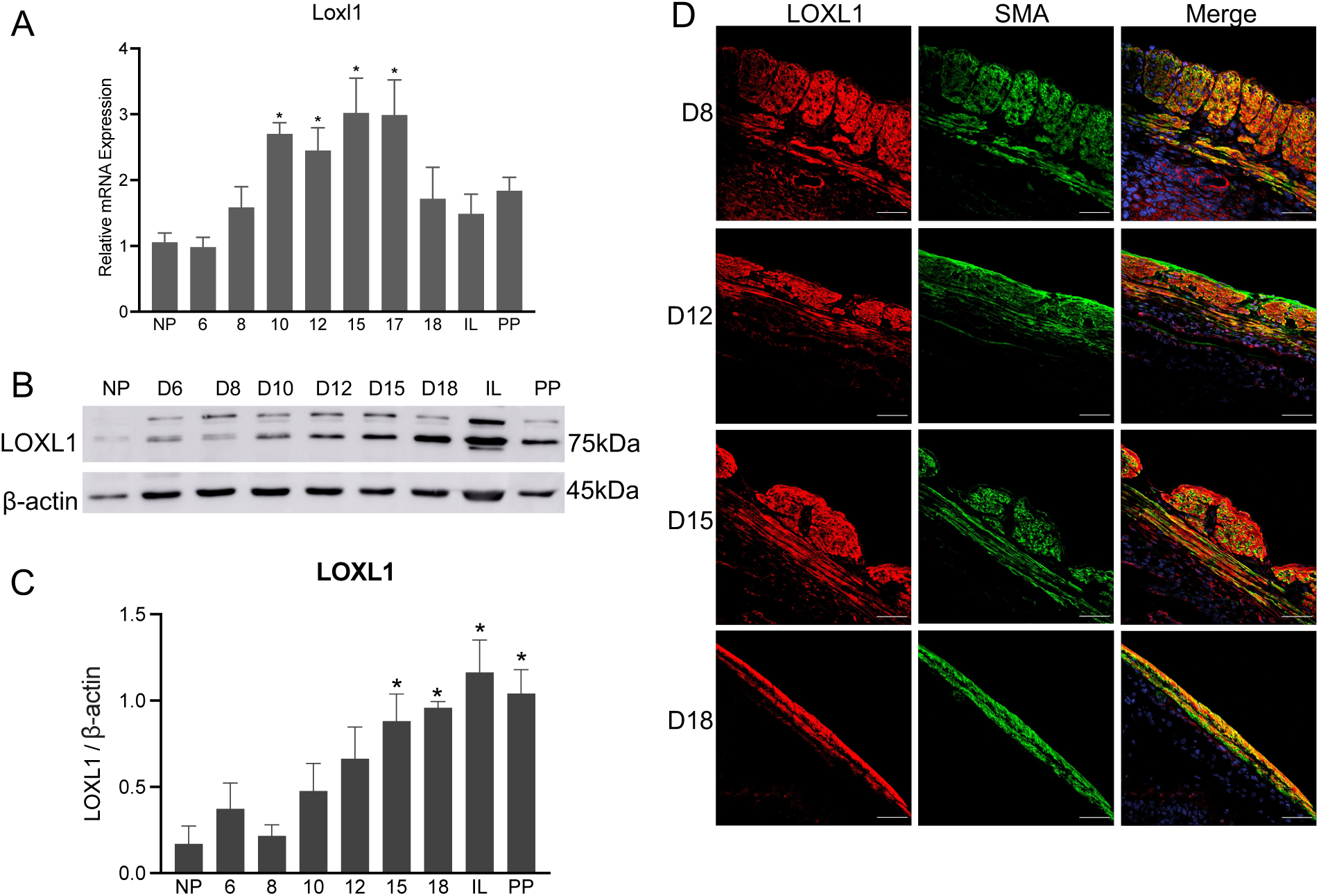
Expression and localization of LOXL1 in mouse myometrium. **A.** Expression of *Loxl1* in mouse myometrium. Total RNA isolated from NP (non-pregnant), gestation day 6, 8, 10, 12, 15, 17, 18, IL (in-labor) and PP (postpartum) mice was used. The mRNA expression was normalized to *Rplp0* and compared with NP samples (n=6/group, *p<0.05). **B.** Western blot analysis of LOXL1 protein levels in myometrial whole tissue lysates of NP, gestation day 6, 8, 10, 12, 15, 18, IL and PP mouse myometrial tissues. β-actin was used as a loading control. These are representative images from three independent replicates. **C.** Quantification of protein levels of LOXL1 isolated from myometrial tissues of NP, gestation day 6, 8, 10, 12, 15, 18, IL and PP mice normalized to β-actin levels and compared with NP samples (n=3/group, *p<0.05). **D.** Confocal imaging analysis of LOXL1 in gestation day 8, 12, 15, and 18 myometrium. Representative images from three independent replicates. Brightness and contrast for each image were optimized individually for optimal visualization of morphology. Scale bar: 50 μm. – LOXL1 Lysyl oxidase like -1, SMA – Smooth muscle actin. Note: LOXL1 is predominantly localized in smooth muscle tissue of the myometrium.

**Fig. 3.**
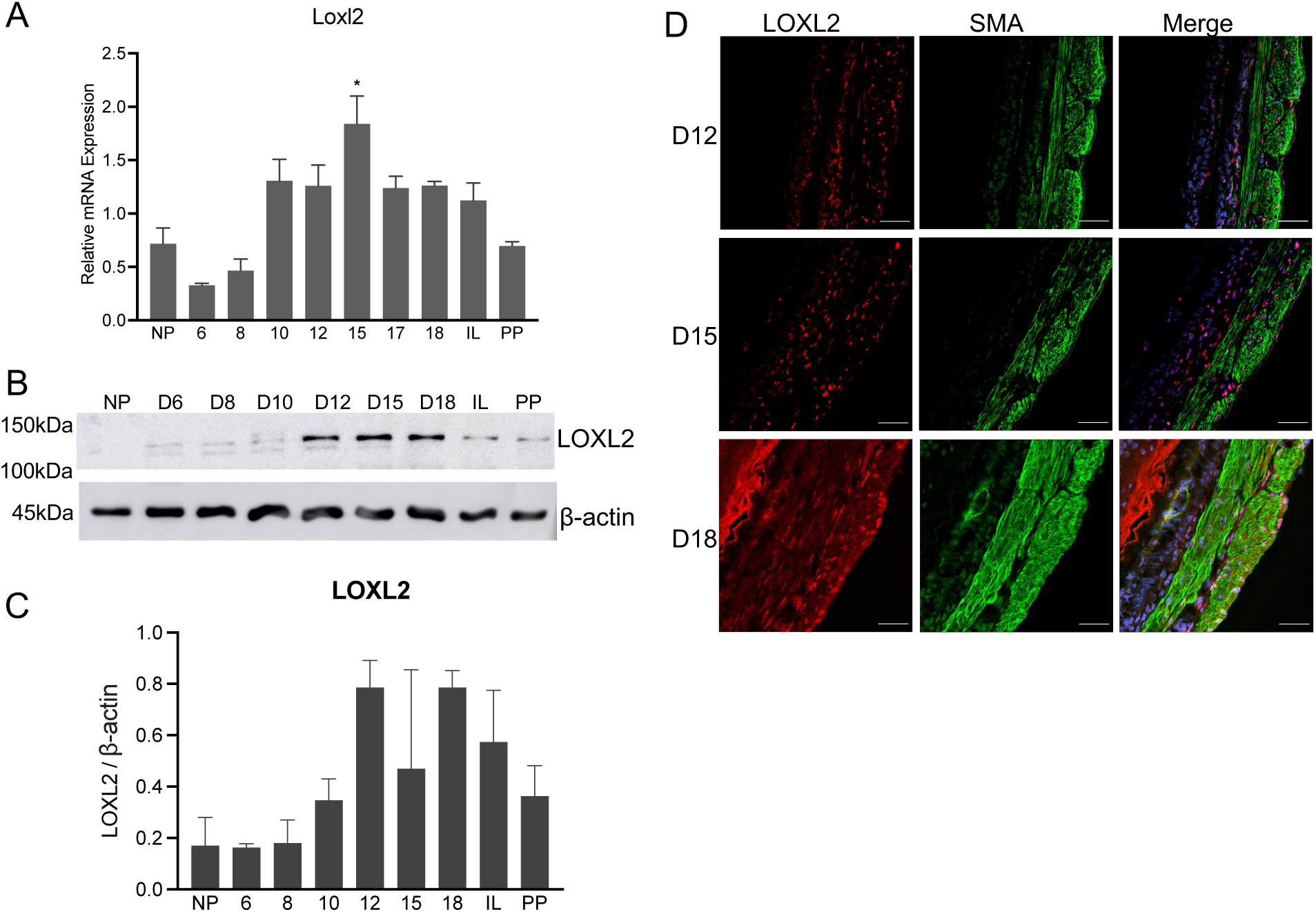
Expression and localization of LOXL2 in mouse myometrium. **A.** Expression of *Loxl2* in mouse myometrium. Total RNA isolated from NP (non-pregnant), gestation day 6, 8, 10, 12, 15, 17, 18, IL (in-labor) and PP (postpartum) mice was used. The mRNA expression was normalized to *Rplp0* and compared with NP samples (n=6/group, *p<0.05). **B.** Western blot analysis of LOXL2 protein levels in myometrial whole tissue lysates of NP, gestation day 6, 8, 10, 12, 15, 18, IL and PP mouse myometrial tissues. β-actin was used as a loading control. These are representative images from three independent replicates. **C.** Quantification of protein levels of LOXL2 isolated from myometrial tissues of NP, gestation day 6, 8, 10, 12, 15, 18, IL and PP mice normalized to β-actin levels and compared with NP samples (n=3/group, *p<0.05). **D.** Confocal imaging analysis of LOXL2 in gestation day 12, 15, and 18 myometrium. Representative images from three independent replicates. Brightness and contrast for each image were optimized individually for optimal visualization of morphology. Scale bar: 50 μm. LOXL2 – Lysyl oxidase like -2, SMA – Smooth muscle actin. Note: LOXL2 is predominantly localized in smooth muscle tissue of the myometrium.

### Enzyme activity of lysyl oxidases increased in pregnant myometrium

Our results so far demonstrated that the lysyl oxidases are significantly induced in the mouse myometrium during pregnancy. The catalytic functions of lysyl oxidases to cross-link collagen and elastin, and thus to modify the structure and function of the ECM, depends on their enzymatic activity. Therefore, we next examined the enzymatic activity of lysyl oxidases in the myometrium using an *in situ* activity assay. We measured the total activity of all lysyl oxidase enzymes because it is not possible to analyze the activity individually. Additionally, this technique is helpful to understand the spatiotemporal activity. The lysyl oxidases were active in all gestational time points as evidenced by signal elicited through the incorporation of EZ-Link Hydrazide-Biotin (BHZ) in the myometrial tissues **(Fig. 4)**. Consistent with the elevated levels of gene and protein levels, the total lysyl oxidase activity was much higher on gestation day 15 compared to all other samples **(Fig. 4)**. These results indicate that the lysyl oxidases are more active in myometrium during late gestation.

**Fig. 4.**
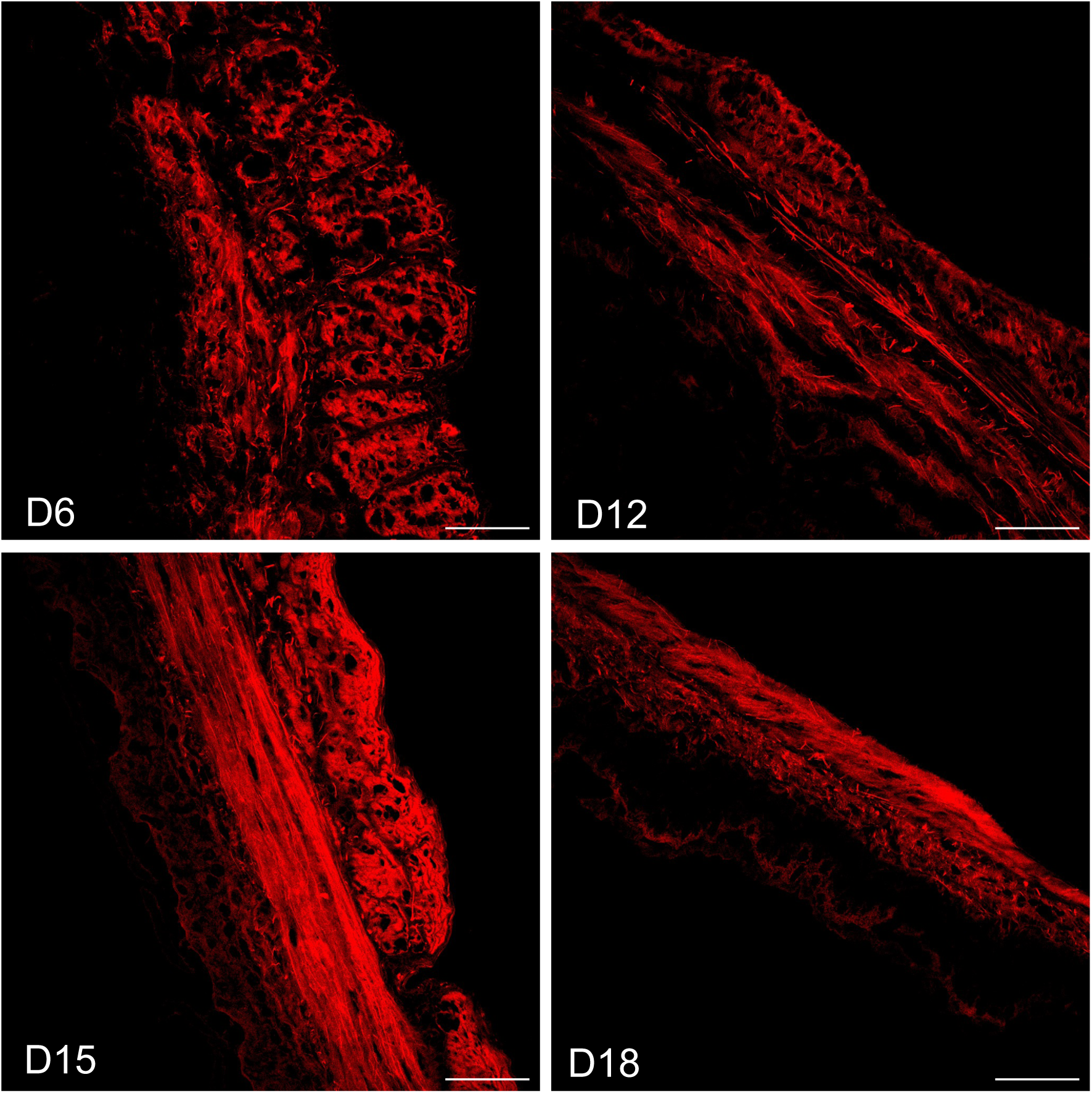
Activity of lysyl xidases in mouse myometrium. Uterine tissue sections prepared from gestation day 6, 12, 15 and 18 mice were incubated with 200 μM biotin-hydrazide (BHZ) overnight and then stained with Alexa Fluor-streptavidin. The representative images from confocal imaging analysis are shown. The intensity red color represents BHZ incorporation (red) which corresponds to activity of the lysyl oxidases. The image settings were set for day 15 sample and used for all other images for comparison. Scale bar: 50 μm.

### Steroid hormones differentially induce Lox and Loxl1-2 in mouse myometrium

The lysyl oxidases were highly induced and active in the pregnant myometrium, indicating possible regulation by steroid hormones. We utilized ovariectomized mice exogenously administered with estrogen, progesterone, or in combination to induce the gene expression of lysyl oxidases in mouse myometrium. In response to 17β-estradiol treatment, *Lox* expression was significantly suppressed, whereas *Loxl1* and *Loxl2* were significantly induced **(Fig. 5A)**. The robust induction of *Loxl2* in response to 17β-estradiol treatment indicates that it could be a potential target of this hormone. Similar to 17β-estradiol, progesterone significantly suppressed the expression of *Lox*. In contrast to 17β-estradiol alone, progesterone or a combination of 17β-estradiol and progesterone did not exert any influence on the expression of *Loxl1* and *Loxl2* **(Fig. 5B)**. It was also possible that 17β-estradiol-induced expression of *Loxl1* and *Loxl2* might have been suppressed by the progesterone. These results indicate that steroid hormones, 17β-estradiol and progesterone, differentially regulate the expression of lysyl oxidases in the myometrium.

**Fig. 5.**
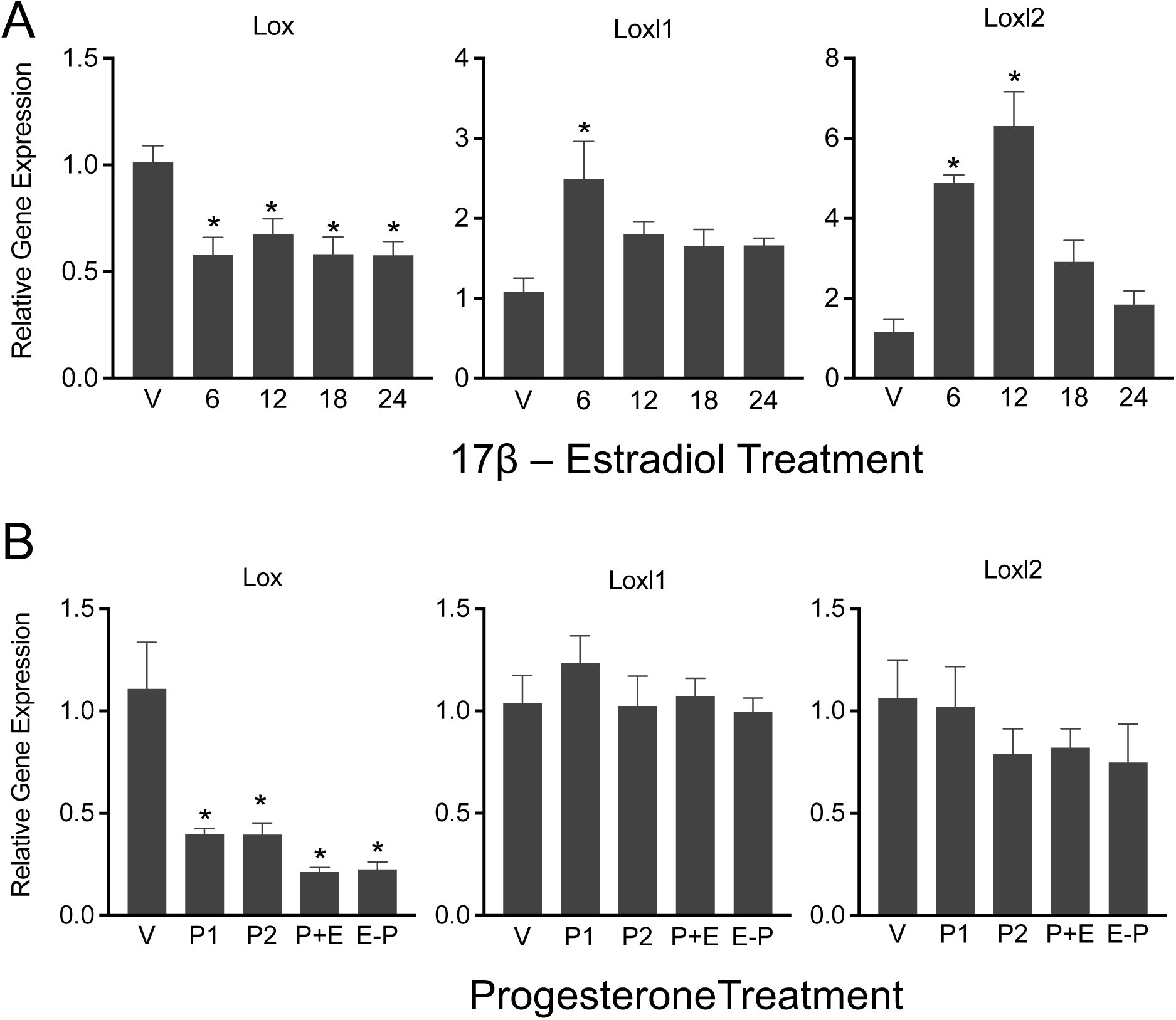
Gene expression of *Lox, Loxl1* and *Loxl2* in response to 17β-estradiol and progesterone in ovariectomized mouse myometrium. **A.** Expression of *Lox, Loxl1* and *Loxl2* in response to 17β-estradiol in ovariectomized mouse myometrium. Ovariectomized mice were treated with vehicle (V), or 17β-estradiol (100 ng/mouse, s/c) for 6, 12, 18 or 24 hours (n=6/group, *p<0.05). **B.** Expression of *Lox, Loxl1* and *Loxl2* in response to progesterone (P) or progesterone and 17β-estradiol (E) in ovariectomized mouse myometrium. Ovariectomized mice were treated with P (1mg/mouse, s/c), P (2mg/mouse, s/c), P (1mg) + E (10ng), E (100ng) – 2 days rest – P (1mg). The tissues were collected after 24 hours (n=6/group, *p<0.05).

### Inhibition of activity of lysyl oxidases delays parturition in mice due to impaired myometrial contractility

To examine the impact of the activity of lysyl oxidases on myometrium and parturition, we treated mice with 0.8% BAPN (A3134, Sigma-Aldrich) through drinking water from gestation day 12 (day of vaginal plug = day 0) and monitored the timing of parturition **(Fig. 6A)**. Lysyl oxidases were induced very early during pregnancy in the myometrium (Fig. 1-3) and were also expressed in multiple other reproductive tissues such placenta, fetal membranes, and the cervix. Using this information, we treated mice from gestation day 12 to avoid the effect of BAPN on placental, fetal membrane and fetal development. In response to BAPN treatment, the parturition process was significantly delayed compared to an untreated control group of mice **(Fig. 6B)**. These results demonstrate that the inhibition of activity of lysyl oxidases delayed parturition in mice. To examine the activity of lysyl oxidases on myometrial contractility, we treated pregnant mice with BAPN from gestation day 12 and collected myometrial tissues on day 18 **(Fig. 6C)**. The expression levels of genes encoding contraction associated proteins – connexin 43 (*Gja1*), oxytocin receptor (*Oxtr*) and prostaglandin endoperoxide synthase 2 (*Ptgs2*) – were significantly reduced in response to BAPN **(Fig. 6D)**. Consistent with mRNA expression, the protein level of connexin 43 was reduced in BAPN treated tissues **(Fig. 6E)**. BAPN treatment did not alter *Col1a1*, *Eln* and *Lox* transcription (Fig 6D). Also, the genes encoding proteins related to myometrial quiescence were not altered in response to BAPN **(Fig. 6F).** These results demonstrate that activity of lysyl oxidases is necessary for myometrial contractility at term to facilitate on-time parturition in mice.

**Fig. 6.**
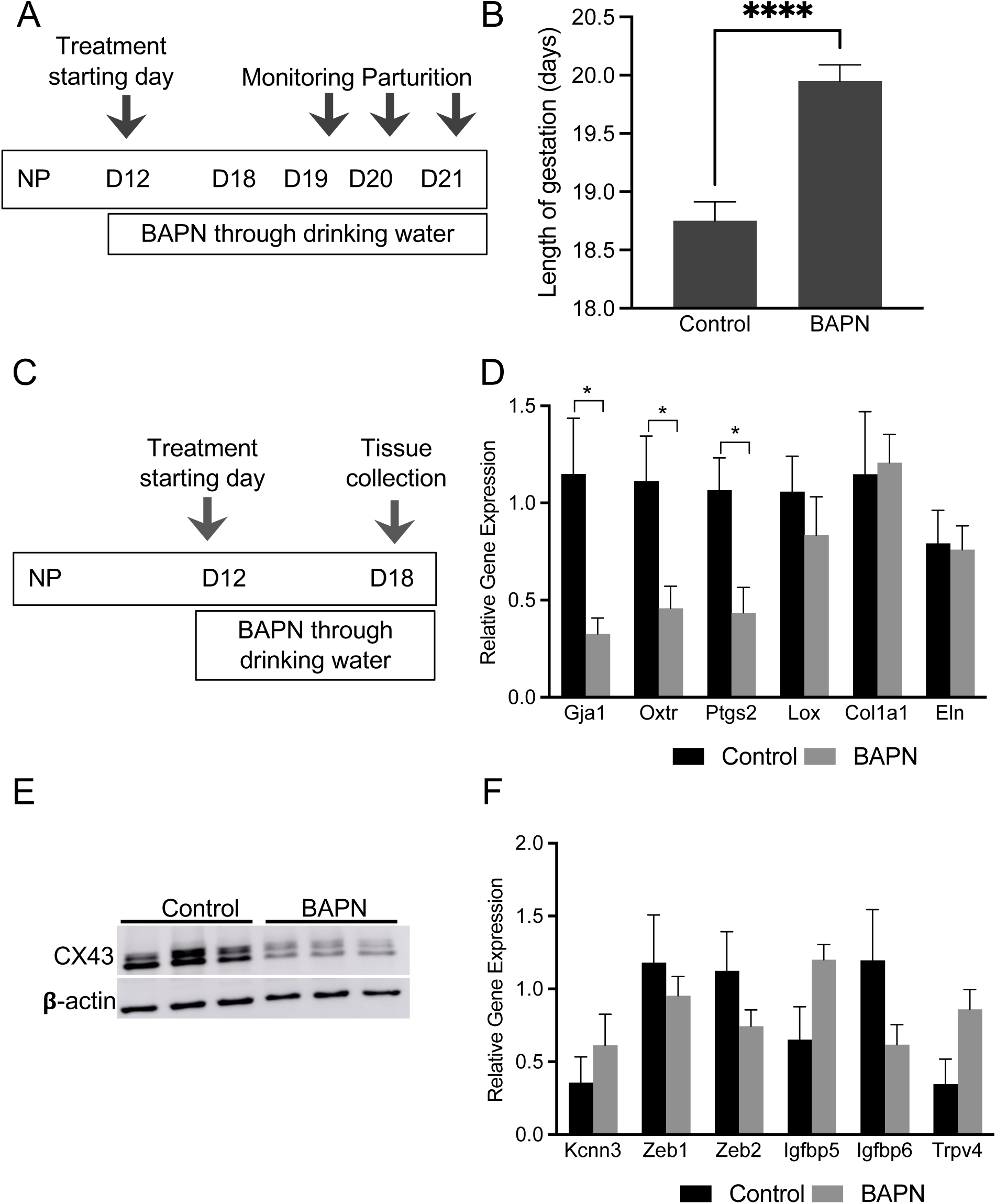
Impact of inhibition of activity of lysyl oxidases in mouse myometrium on parturition-timing and myometrial contractility. **A.** Schematic representation of BAPN treatment to monitor parturition timing in mice. Pregnant mice were treated with BAPN through drinking water from gestation day 12 until parturition. **B.** The parturition-timings between control (n = 10) and BAPN treated (n = 8) mice were monitored, recorded and analyzed. In BAPN treated group, the parturition was significantly delayed compared to the control group (*p<0.05). **C.** Schematic representation of BAPN treatment for tissue collection. Pregnant mice were treated with BAPN through drinking water from gestation day 12 until gestation day 18 for the collection of myometrial tissues to analyze gene and protein expression. Untreated mice served as control (n=6/group). **D.** Expression of connexin 43 (*Gja1*), oxytocin receptor (*Oxtr*), prostaglandin endoperoxide synthase 2 (*Ptgs2*), lysyl oxidase (*Lox),* collagen type I alpha 1 chain (*Col1a1*) and elastin (*Eln*) in the myometrial tissues collected on gestation day 18 from control and BAPN treated mice (n=6/group). Expression of genes encoding contraction associated proteins – *Gja1*, *Oxtr*, and *Ptgs2* – are significantly reduced in response to BAPN treatment (*p<0.05). The expression levels of *Lox, Col1a1* and *Eln* are not altered in response to BAPN. **E.** Western blot analysis of protein levels of connexin 43 (39–44kDa) in control and BAPN treated myometrial tissues collected on gestation day 18 (n=3/group). β-actin was used as a loading control. The levels of connexin 43 are reduced in response to BAPN indicating myometrial contractile dysfunction. **F.** Expression of *Kcnn3, Zeb1* and *2*, *Igfbp5* and *6*, and *Trvp4* – genes encoding factors known to be involved in the maintenance of uterine quiescence – are not altered in response to BAPN treatment. The total RNA isolated from myometrial tissues collected on gestation day 18 from control and BAPN treated mice were used (n=6/group).

### Lysyl oxidases maintain integrity of the ECM in myometrium during pregnancy and parturition

To examine the activity of lysyl oxidases on the integrity of myometrial ECM, we compared the proteomic profiles of myometrial tissues isolated from control and BAPN-treated mice using isobaric tandem mass tags, a stable isotope-based quantitative proteomic approach **(Fig 7A).** A total of 5,176 proteins were quantified and represented in a volcano plot (**Fig. 7B)**. 356 proteins were identified as differently expressed in BAPN vs. control (two-tailed t-test; *p* < 0.05) (**Fig. 7B)**. 22 differentially expressed proteins had a 2-fold increase (20) or decrease (2) in abundance in the BAPN-treated samples are represented in a heat map (**Fig. 7C)** and also listed in **Table 2**. Term enrichment analysis on the 22 differentially expressed proteins indicated that collagens are involved in several pathways **(Fig. 7D** and **Supplementary table).** The top 3 terms are protein digestion and absorption, ECM-receptor interaction, and focal adhesion **(Fig. 7E).** Other differentially expressed proteins include peroxidasin homolog (*Pxdn*), hemoglobin subunit epsilon-Y2 (*Hbb-y*), leucine-rich HEV glycoprotein (*Lrg1*), hemoglobin subunit zeta (*Hbz*) serum amyloid P-component (*Apcs*), alpha-1-acid glycoprotein 1 (*Orm1*), microfibril-associated glycoprotein 1 (*Mfap2*), sushi-repeat-containing protein (*Srpx*), aldo-keto reductase family 1 member C18 (*Akr1c18*), aquaporin-5 (*Aqp5*), OCIA domain-containing protein 2 (*Ociad2*), transforming growth factor beta (*Tgfb2*), alpha-fetoprotein (*Afp*), and microfibril-associated glycoprotein 4 (*Mfap4*). Consistent with gene expression, the protein level of Gap junction alpha-1 protein (*Gja1*) was decreased in BAPN treated samples. Of the 5176 quantified proteins, 131 were labeled as ECM-related according to Gene Ontology (Cellular Component). 96 of which were classified in the matrisomeDB (https://matrisomedb.org/) (38) into categories under two divisions (Core matrisome and Matrisome-associated) **(Fig. 8A** and **Table 3).** ECM proteins that were quantified to have differential abundances (p < 0.05) in the BAPN samples are represented in heatmaps **(Fig. 8B)** and are also listed in **Table 4**. Majority of the differentially expressed (p <0.05) ECM proteins were found to be upregulated. Elastin and LOXL1 were significantly increased in the BAPN-samples with a 1.6-fold upregulation. These results revealed that proteomic composition of myometrial ECM is altered upon inhibition of the activity of lysyl oxidase enzymes.

**Fig. 7.**
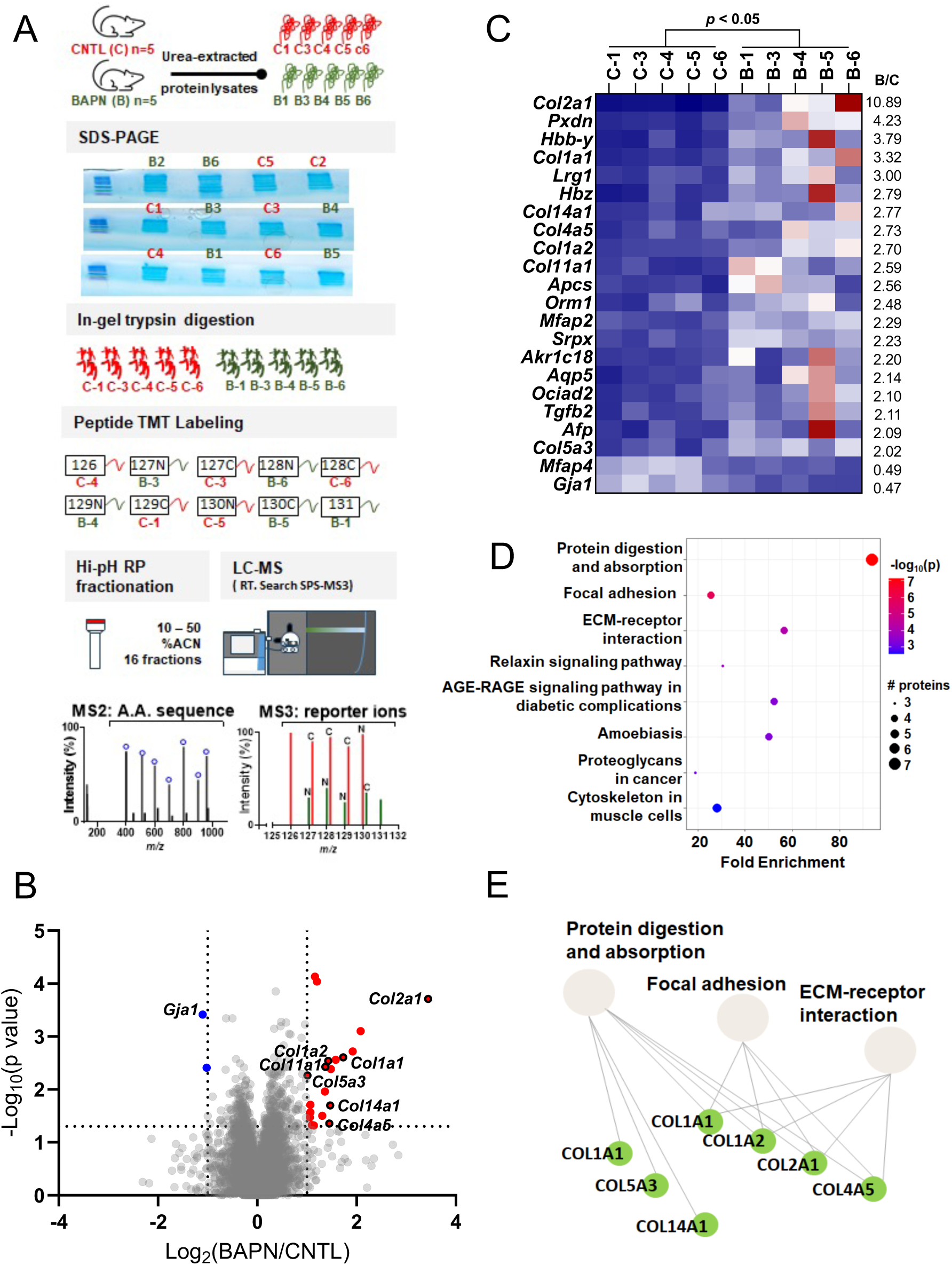
Proteomic analysis reveals alterations in the composition of myometrial proteome between control and BAPN treated mice. **A.** Experimental design. Urea-extracted protein samples from myometrial tissues of control and BAPN treated mice (n = 6/group) were loaded onto SDS-PAGE slightly into the separating gel. Five samples per group were subsequently used. After in-gel digestion, peptide labeling with tandem mass tags (TMT), and high-pH reversed-phase (RP) fractionation, peptides were identified in tandem mass spectrometry (MS2) and their reporter ions quantified in MS3 scans after Real time search (RTS) and Synchronous Precursor Selection (SPS). **B.** Volcano plot. 5,176 proteins were quantified and represented in a volcano plot. 356 proteins were identified as differently expressed in BAPN vs. control (two-tailed t-test; *p* < 0.05) with 20 and 2 proteins having a 2-fold increase or decrease in abundance (BAPN/Control: log2 < -1 or >1), respectively. Differentially expressed proteins are highlighted (upregulated in red; downregulated in blue) Collagens and Gja1 are labeled with their gene names. Cut-offs of fold change at 2 (log2 = -1 or 1) and *p*-value at 0.05 (-log10 0.05 = 1.301) are indicated by dotted lines on the x and *y-*axis, respectively. **C.** Heat maps of the differential proteomes. Scaled abundances of the 22 differentially expressed proteins are represented in a heat map, listed (top to bottom) according to their average log2 fold change from high to low. The scale of the color intensity is arbitrary (double gradient). **D.** Enrichment Analysis. Top 8 enrichment terms of the differentially expressed proteins (BAPN/Control > 2, p < 0.05). **E.** Interactions between the top 3 enrichment terms. Upregulated proteins are highlighted in green.

**Fig. 8.**
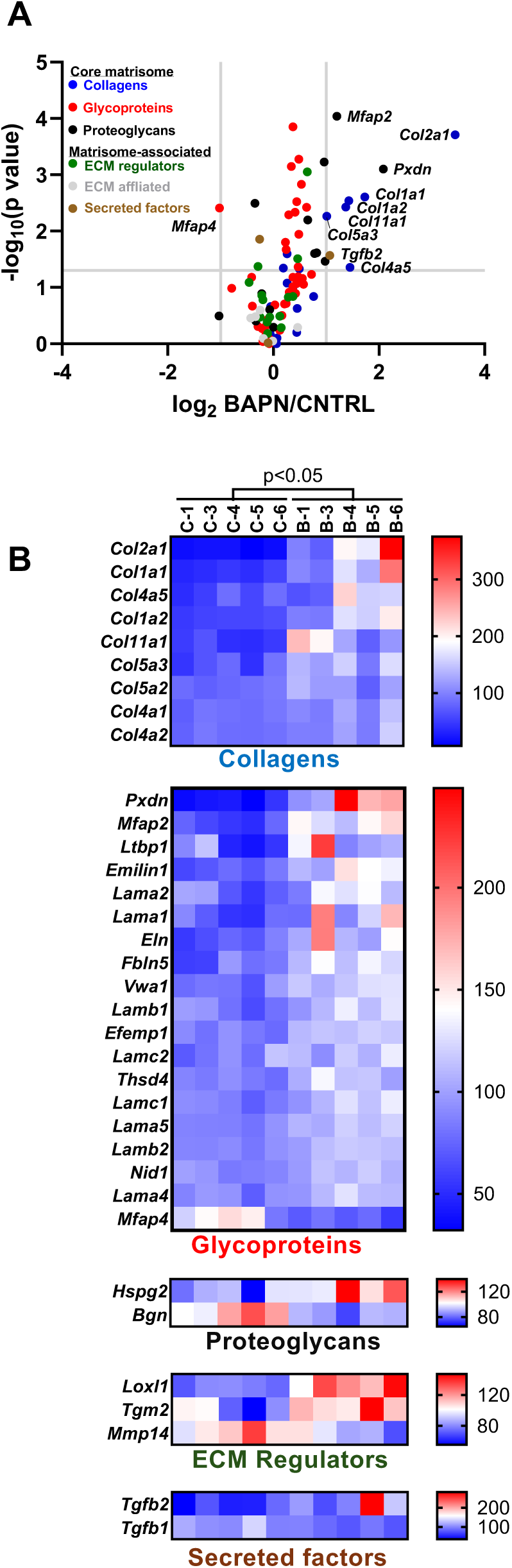
Myometrial extracellular matrix (ECM) proteome in BAPN-treated mice. **A.** Volcano plot. ECM proteins were colored according to the categories they belong to and those that were quantified with a fold change (BAPN/Control) >2 and p <0.05 were labeled. Cut-offs of fold change at 2 (log2 = -1 or 1) and *p*-value at 0.05 (-log10 0.05 = 1.301) are indicated by dotted lines on the x and *y-*axis, respectively. **B.** Heat map of the differentially expressed ECM proteins (p <0.05) of various ECM categories. Scaled abundances of the differentially expressed proteins are represented and listed (top to bottom) according to their average log2 fold change from large to small. The scale of the color intensity is arbitrary (double gradient).

**Table 2.** Differential proteome of the myometrium in BAPN-treated mice.

| Accession | Gene Symbol | Description | Fold Change<br>(BAPN/CNTL) | P<br>(**<0.05**) |
| --- | --- | --- | --- | --- |
| P28481 | *Col2a1* | Collagen alpha-1(II) chain | 10.89 | 1.94E-04 |
| Q3UQ28 | *Pxdn* | Peroxidasin homolog | 4.23 | 7.90E-04 |
| P02104 | *Hbb-y* | Hemoglobin subunit epsilon-Y2 | 3.79 | 1.90E-03 |
| P11087 | *Col1a1* | Collagen alpha-1(I) chain | 3.32 | 2.49E-03 |
| Q91XL1 | *Lrg1* | Leucine-rich HEV glycoprotein | 3.00 | 2.76E-03 |
| P06467 | *Hbz* | Hemoglobin subunit zeta | 2.79 | 4.13E-03 |
| B7ZNH7 | *Col14a1* | Collagen alpha-1(XIV) chain | 2.77 | 2.01E-02 |
| Q63ZW6 | *Col4a5* | Col4a5 protein | 2.73 | 4.42E-02 |
| Q01149 | *Col1a2* | Collagen alpha-2(I) chain | 2.70 | 2.90E-03 |
| Q61245 | *Col11a1* | Collagen alpha-1(XI) chain | 2.59 | 3.74E-03 |
| P12246 | *Apcs* | Serum amyloid P-component | 2.56 | 1.09E-02 |
| Q60590 | *Orm1* | Alpha-1-acid glycoprotein 1 | 2.48 | 3.16E-02 |
| Q99PM0 | *Mfap2* | Microfibril-associated glycoprotein 1 | 2.29 | 9.10E-05 |
| Q9R0M3 | *Srpx* | Sushi-repeat-containing protein SRPX | 2.23 | 7.39E-05 |
| Q8K023 | *Akr1c18* | Aldo-keto reductase family 1 member C18 | 2.20 | 4.79E-02 |
| Q9WTY4 | *Aqp5* | Aquaporin-5 | 2.14 | 4.70E-02 |
| Q9D8W7 | *Ociad2* | OCIA domain-containing protein 2 | 2.10 | 1.94E-02 |
| A0A0A6YXU9 | *Tgfb2* | Transforming growth factor beta | 2.11 | 2.70E-02 |
| P02772 | *Afp* | Alpha-fetoprotein | 2.09 | 3.36E-02 |
| Q9JLI2 | *Col5a3* | Collagen type V alpha 3 chain | 2.02 | 5.42E-03 |
| Q9D1H9 | *Mfap4* | Microfibril-associated glycoprotein 4 | 0.49 | 3.87E-03 |
| P23242 | *Gja1* | Gap junction alpha-1 protein | 0.47 | 3.84E-04 |

**Table 3.**
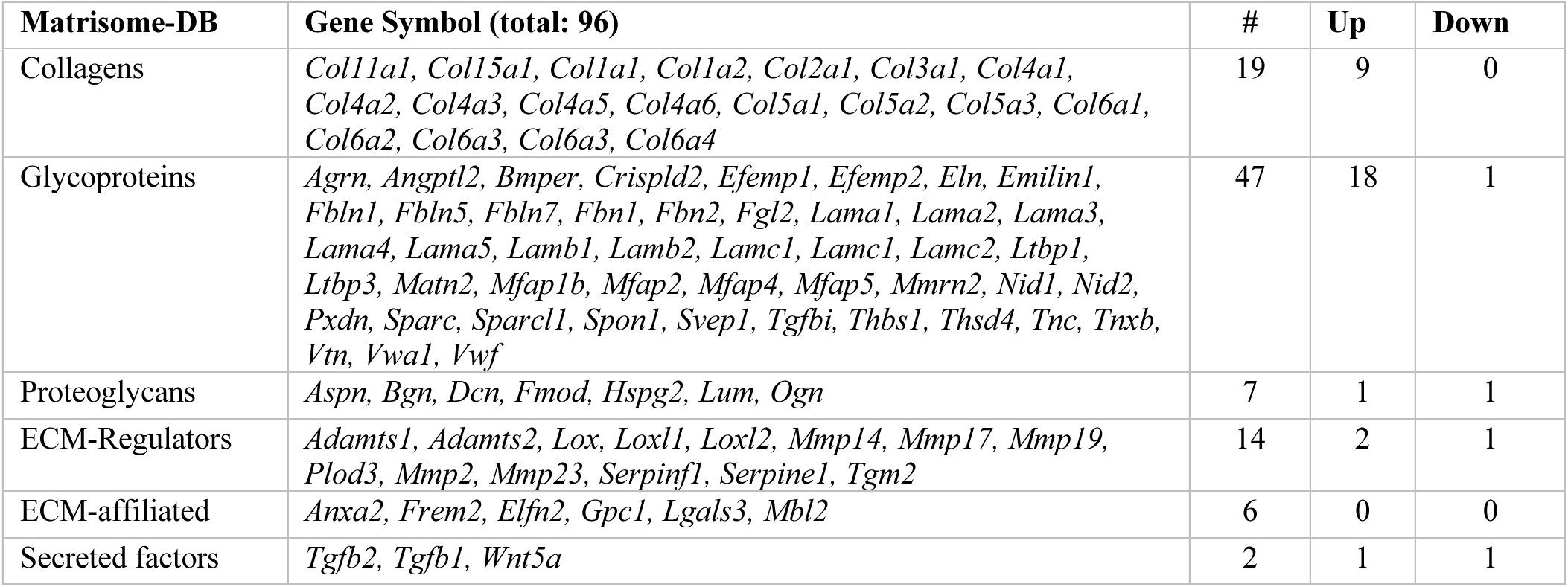
Quantified extracellular matrix (ECM) proteome of the myometrium in BAPN-treated mice.

**Table 4.**
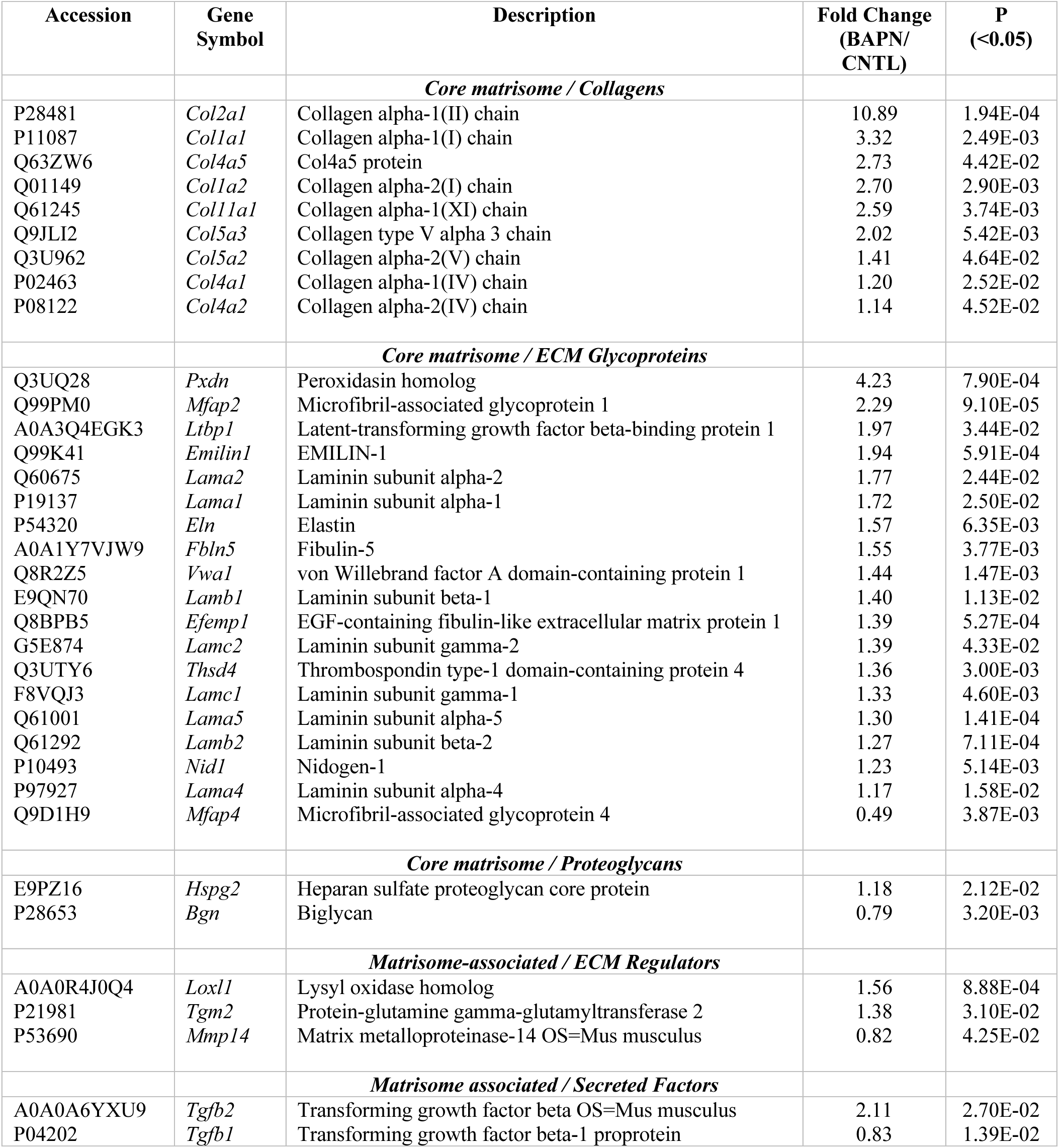
Differential extracellular matrix (ECM) proteome of the myometrium in BAPN-treated mice.

To further validate our results obtained from the proteomic profiling, we performed western blot analysis. The myometrial tissues isolated on gestation day 18 from control and BAPN treated mice were subjected to PBS and followed by urea extraction to analyze the protein levels of collagen I, III and elastin. The protein levels of collagen I, III and elastin were higher in the BAPN treated group compared to control **(Fig. 9).** This result demonstrates that inhibition of LOX activity presumably leads to less crosslinked and weaker collagen and elastin which makes them easily extractable. Collectively, these results demonstrate that the lysyl oxidases maintain integrity of the ECM in myometrium during pregnancy.

**Fig. 9.**
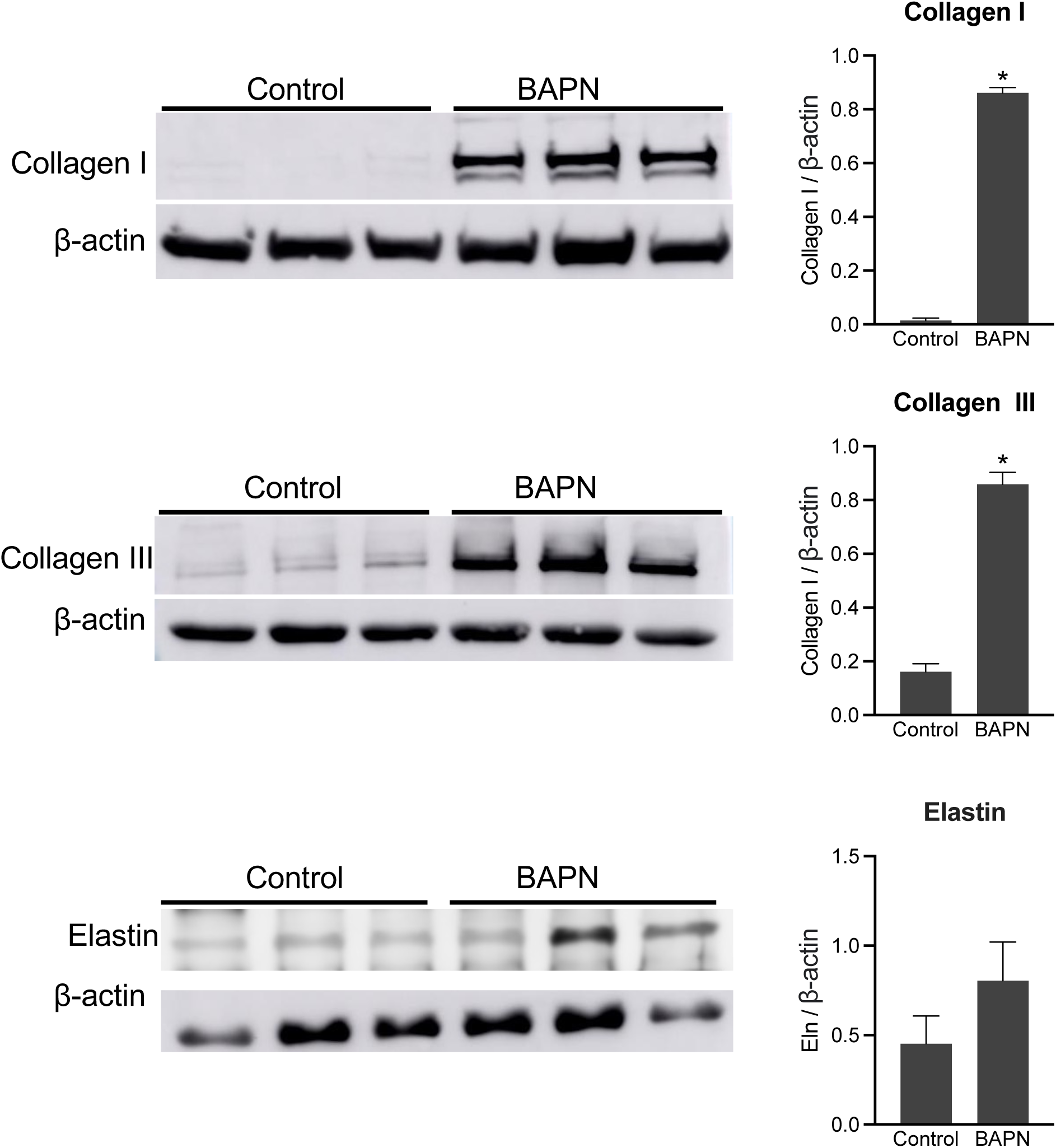
Impact of inhibition of activity of lysyl oxidases on the integrity of myometrial extracellular matrix. Western blot analysis of collagen I (140–210kDa), collagen III (140– 180kDa), and tropoelastin (monomer: 72kDa; dimer:144kDa) in urea fractions of myometrial tissues collected from gestation day 18 mice (left panel). Pregnant mice were treated with BAPN through drinking water from gestation day 12 until gestation day 18. Untreated mice served as control (n=3/group). Quantification of levels of Collagen I, III and elastin in BAPN treated samples compared with control samples (right panel). The protein levels were normalized to β-actin levels (n=3/group, *p<0.05). Note: The protein levels of collagen I, III and elastin were higher in BAPN treated group compared to untreated control group because these extracellular matrix proteins were easily extractable in BAPN treated tissues.

## DISCUSSION

This study expands our understanding of myometrial tissue function beyond smooth muscle cells to include ECM to better outline the myometrial transition from quiescence to contractility at parturition. Though the understanding of the unequivocal role of smooth muscle cells in myometrial function has been demonstrated, contribution of myometrial ECM in the regulation of myometrial function is an understudied area. The predominant ECM components such as fibrillar collagen and elastic fibers undergo structural remodeling in the mouse myometrium during pregnancy (15). The synthesis, processing and assembly of these fibers are regulated by multiple factors involving a complex multistep process. Among all factors, lysyl oxidase enzymes are emerging as critical targets because of their ability to crosslink collagen and elastic fibers and thus mediate the strength and stability of the ECM. The current study provides compelling evidence that lysyl oxidases are critical mediators of myometrial ECM integrity to maintain myometrial tissue function during pregnancy and parturition. The results of this study demonstrate that a subset of these lysyl oxidase enzymes is highly induced in the pregnant mouse myometrium. The inhibition of activity of these enzymes results in delayed parturition due to the failure of myometrium to mount a contractile response at an appropriate time to initiate parturition. Being known modifiers of the ECM structure and function, lysyl oxidases play a critical role in myometrial tissue mechanical function by maintaining structural integrity of collagen and elastic fibers as well as composition of myometrial ECM. In this study, we have uncovered a novel role of lysyl oxidases mediating myometrial contractility for parturition timing in mice.

The ECM remodeling is unique to each and every compartment of the female reproductive tract in alignment with their corresponding physiological function. For example, ECM remodeling with respect to fibrillar collagen and elastic fiber reorganization in the cervix and myometrium has unique differences to accomplish their function during pregnancy and parturition (15,34,39,40). In myometrium, thin and scattered collagen fibers are reorganized into thick and straight bundles as pregnancy progresses, whereas in the cervix these collagen fibers are reorganized from thin and linear fibers to thick and wavy bundles (15,34). These structural alterations support their respective functions in the myometrium and cervix to accommodate fetal growth and to undergo softening to prepare for birth, respectively. These reorganizational changes to structure and function are mediated by new synthesis as well as differential expression of factors involved in the synthesis, processing and assembly of collagen and elastic fibers. Notably, the cross-linking lysyl oxidase enzymes which determine strength and stability of collagen and elastic fibers are differentially expressed in myometrium and cervix during pregnancy. In mouse cervix, the level of LOX transcript and activity are high during NP, decline early in pregnancy and remain low until gestation day 18 (34). This temporal regulation has direct functional implications in the cervix. In the non-pregnant mouse cervix, the high level of LOX increases the cross-links of collagen and elastic fibers to maintain mechanical strength. In early pregnancy LOX activity declines in the cervix, leading to a reduction in the level of cross-links, change in the ECM structure and a decline in mechanical stiffness which subsequently leads to progressive softening (33,34,41). In mouse myometrium, Lox transcripts are significantly induced in the quiescent phase of myometrial remodeling similar to the temporal expression of collagen and elastin. These elevated activities presumably increase collagen and elastin cross-links, promote change in the ECM structure and increase mechanical stiffness of the myometrium during pregnancy to support mechanical loading exerted by the growing fetus and also to mount forceful contractile response at term for the expulsion of the fetus. Ongoing functional and mechanistic studies in our laboratory are aimed to test these hypotheses in the myometrium.

In most tissues of the body, the ECM is laid during embryonic development and undergoes limited remodeling in the adult life (42–44). Unlike other adult tissues, ECM present in the reproductive tissues undergo progressive remodeling, followed by constant turn over to meet the demands exerted by the cyclical processes of pregnant, parturient and non-pregnant states of female’s lives (18,34,45,46). Because of these dynamic changes, the synthesis of ECM factors in the reproductive tissues is under the control of steroid hormones, estrogen and progesterone, unlike most adult tissues (15,16,34). Our previous and current studies indicate that many genes encoding ECM proteins are differentially regulated by steroid hormones in mouse myometrium. Comparison of the expression pattern of LOX in the cervix and myometrium reveals that LOX is differentially regulated in the same steroid hormonal milieu in distinct reproductive tissues to achieve different functions. These novel findings suggest that the steroid hormonal regulation of lysyl oxidases in the myometrium is further modified by cell type specific regulation potentially by other factors such as HIF, TGFβ, microRNAs, small leucine rich proteoglycans or fibulins.

Premature myometrial contractility is a notable phenomenon in preterm birth. Therefore, understanding the mechanisms involved in the transition of myometrial quiescence to contractility will provide opportunities to develop therapeutic strategies to clinically manage preterm birth. Our understanding on the pathways regulating myometrial contractility such as hormonal, inflammatory and fetal signaling pathways have improved over the years but the mechanical signaling involving ECM is very limited. Understanding the role of components of the ECM might help to uncover potential targets for the development of therapeutic strategies to clinically manage premature myometrial contractility as well as problems associated with labor at birth. The function of lysyl oxidases has received wide attention in pathological conditions such as fibrosis and tumor development. In these pathological conditions, aberrant deposition of the ECM, particularly collagen and tissue rigidity invariably result in loss of cell and tissue function. Most of these pathological conditions are also characterized by elevated synthesis and activity of lysyl oxidases. Therefore, the development of treatment strategies targeting lysyl oxidases to inhibit their activity has become an attractive arena for therapy (19,47).

This study has a few limitations. The lysyl oxidases are highly induced very early in pregnancy. However, BAPN treatment was administered on gestation day 12 to avoid its impact on the fetus, fetal membrane and placental development. Therefore, complete inhibition of lysyl oxidases in myometrium might not be possible through this approach. As previously mentioned, lysyl oxidases are widely expressed in almost all reproductive and fetal tissues. Therefore, the results of this study might have been confounded by widespread effect of BAPN on all of these tissues. In addition, there are five lysyl oxidases and three of them are highly induced in pregnant mouse myometrium. Because of their functional redundancy, identifying their unique function might be very challenging. To circumvent all these confounding factors, the lysyl oxidases should be specifically deleted in the myometrium to understand their distinct function in the myometrium during pregnancy and parturition. Despite these limitations, this study has provided important information on the function of lysyl oxidases in regulation of the structure and function of myometrial ECM and their impact in the processes of pregnancy and parturition.

## DATA AVAILABILITY

Original data generated and analyzed during this study are included in this published article or in supplementary materials. The mass spectrometry proteomics data have been deposited to the ProteomeXchange Consortium via the PRIDE partner repository with the dataset identifier PXD055537.

## ACKNOWLEDGEMENTS

This work was supported by funding from the National Institutes of Health K99/R00HD090301(SN). The content is solely the responsibility of the authors and does not necessarily represent the official views of the National Institutes of Health. Confocal microscopy was performed on a Nikon A1R-HD point scanning confocal supported by NIH award number 1S10OD025030-01 from the Office of Research Infrastructure Programs. We sincerely acknowledge Dr. Douglas Taatjes, Facility Director and Nicole Bouffard, staff at the Microscopy Imaging Center at the University of Vermont (RRID# SCR_018821) for their help and support. The Vermont Biomedical Research Network Proteomics Facility (RRID: SCR_018667) is supported through NIH grant P20GM103449 from the INBRE Program of the National Institute of General Medical Sciences.

## AUTHOR CONTRIBUTIONS

SN conceptualized the study, designed the experiments, and wrote the manuscript. SN, AO, CD, SG, and YL conducted experiments, and analyzed the data. SN, AO, CD, SG, MM and YL reviewed, edited and approved the manuscript.

