## Supplementary table 1 for "Lysyl Oxidases are Necessary for Myometrial Contractility and On-Time Parturition in Mice"

### pathfindR - All Enriched Terms

| ID | Term_Description | Fold_Enrichment | occurrence | support | lowest_p | highest_p | Up_regulated | Down_regulated |
|---|---|---|---|---|---|---|---|---|
| hsa04974 | Protein digestion and absorption | 80.875383 | 10 | 0.2728682 | 7.4e-08 | 3.7e-07 | COL1A1, COL1A2, COL2A1, COL4A5, COL11A1, COL5A3, COL14A1 | |
| hsa04510 | Focal adhesion | 22.113392 | 10 | 0.0909561 | 1.5e-06 | 7.3e-06 | COL1A1, COL1A2, COL2A1, COL4A5 | |
| hsa04512 | ECM-receptor interaction | 48.965368 | 10 | 0.1830233 | 6.5e-05 | 1.6e-04 | COL1A1, COL1A2, COL2A1, COL4A5 | |
| hsa04926 | Relaxin signaling pathway | 26.365967 | 9 | 0.0222222 | 1.8e-04 | 1.8e-04 | COL1A1, COL1A2, COL4A5 | |
| hsa04933 | AGE-RAGE signaling pathway in diabetic complications | 45.198801 | 10 | 0.1522459 | 4.1e-04 | 1.6e-02 | TGFB2, COL1A1, COL1A2, COL4A5 | |
| hsa05146 | Amoebiasis | 43.295694 | 10 | 0.1522459 | 4.7e-04 | 1.7e-02 | TGFB2, COL1A1, COL1A2, COL4A5 | |
| hsa05205 | Proteoglycans in cancer | 16.235885 | 9 | 0.0222222 | 7.7e-04 | 7.7e-04 | TGFB2, COL1A1, COL1A2 | |
| hsa04820 | Cytoskeleton in muscle cells | 24.137857 | 8 | 0.0232558 | 2.7e-03 | 2.7e-03 | COL1A1, COL1A2, COL11A1, COL5A3, COL4A5 | |
| hsa05144 | Malaria | 23.913319 | 9 | 0.0222222 | 3.5e-03 | 3.5e-03 | TGFB2 | |
| hsa05321 | Inflammatory bowel disease | 17.728840 | 9 | 0.0222222 | 6.4e-03 | 6.4e-03 | TGFB2 | |
| hsa05211 | Renal cell carcinoma | 16.585044 | 9 | 0.0222222 | 7.3e-03 | 7.3e-03 | TGFB2 | |
| hsa05140 | Leishmaniasis | 15.121658 | 9 | 0.0222222 | 8.9e-03 | 8.9e-03 | TGFB2 | |
| hsa05212 | Pancreatic cancer | 14.689610 | 9 | 0.0222222 | 9.4e-03 | 9.4e-03 | TGFB2 | |
| hsa05220 | Chronic myeloid leukemia | 14.281566 | 9 | 0.0222222 | 9.9e-03 | 9.9e-03 | TGFB2 | |
| hsa04540 | Gap junction | 13.895577 | 9 | 0.0232558 | 1.0e-02 | 2.1e-02 | | GJA1 |
| hsa05210 | Colorectal cancer | 12.853409 | 9 | 0.0222222 | 1.2e-02 | 1.2e-02 | TGFB2 | |
| hsa05323 | Rheumatoid arthritis | 12.694725 | 9 | 0.0222222 | 1.3e-02 | 1.3e-02 | TGFB2 | |
| hsa05410 | Hypertrophic cardiomyopathy | 11.425252 | 9 | 0.0222222 | 1.6e-02 | 1.6e-02 | TGFB2 | |
| hsa05145 | Toxoplasmosis | 10.711174 | 9 | 0.0222222 | 1.8e-02 | 1.8e-02 | TGFB2 | |
| hsa05414 | Dilated cardiomyopathy | 10.711174 | 9 | 0.0222222 | 1.8e-02 | 1.8e-02 | TGFB2 | |
| hsa05142 | Chagas disease | 10.600750 | 9 | 0.0222222 | 1.8e-02 | 1.8e-02 | TGFB2 | |
| hsa04350 | TGF-beta signaling pathway | 10.282727 | 9 | 0.0232558 | 1.9e-02 | 1.9e-02 | TGFB2 | |
| hsa04068 | FoxO signaling pathway | 8.498122 | 9 | 0.0222222 | 2.8e-02 | 2.8e-02 | TGFB2 | |
| hsa04380 | Osteoclast differentiation | 8.292522 | 9 | 0.0222222 | 3.0e-02 | 3.0e-02 | TGFB2 | |
| hsa05226 | Gastric cancer | 7.292714 | 9 | 0.0222222 | 3.8e-02 | 3.8e-02 | TGFB2 | |
| hsa04110 | Cell cycle | 7.091536 | 9 | 0.0222222 | 4.0e-02 | 4.0e-02 | TGFB2 | |
| hsa04218 | Cellular senescence | 6.995053 | 9 | 0.0222222 | 4.2e-02 | 4.2e-02 | TGFB2 | |
| hsa05161 | Hepatitis B | 6.901159 | 9 | 0.0222222 | 4.3e-02 | 4.3e-02 | TGFB2 | |
| hsa04390 | Hippo signaling pathway | 13.710303 | 9 | 0.0222222 | 4.3e-02 | 4.3e-02 | TGFB2, AFP | |
| hsa05225 | Hepatocellular carcinoma | 6.549508 | 9 | 0.0222222 | 4.7e-02 | 4.7e-02 | TGFB2 | |
